# Explosive sensing with insect-based biorobots

**DOI:** 10.1101/2020.02.10.940866

**Authors:** Debajit Saha, Darshit Mehta, Ege Atlan, Rishabh Chandak, Mike Traner, Ray Lo, Prashant Gupta, Srikanth Singamaneni, Shantanu Chakrabartty, Barani Raman

**Affiliations:** Department of Biomedical Engineering, Washington University in St. Louis; Department of Electrical and Systems Engineering, Washington University in St. Louis; Department of Mechanical Engineering and Material Science, Washington University in St. Louis

## Abstract

Stand-off chemical sensing is an important capability with applications in several domains including homeland security. Engineered devices for this task, popularly referred to as electronic noses, have limited capacity compared to the broad-spectrum abilities of the biological olfactory system. Therefore, we propose a hybrid bio-electronic solution that directly takes advantage of the rich repertoire of olfactory sensors and sophisticated neural computational framework available in an insect olfactory system. We show that select subsets of neurons in the locust (*Schistocerca americana*) brain were activated upon exposure to various explosive chemical species (such as DNT and TNT). Responses from an ensemble of neurons provided a unique, multivariate fingerprint that allowed discrimination of explosive vapors from non-explosive chemical species and from each other. Notably, target chemical recognition could be achieved within a few hundred milliseconds of exposure. Finally, we developed a minimally-invasive surgical approach and mobile multi-unit electrophysiological recording system to tap into the neural signals in a locust brain and realize a biorobotic explosive sensing system. In sum, our study provides the first demonstration of how biological olfactory systems (sensors and computations) can be hijacked to develop a cyborg chemical sensing approach.

**SUMMARY:** We demonstrate a bio-robotic chemical sensing approach where signals from an insect brain are directly utilized to detect and distinguish various explosive chemical vapors.

## Introduction

Rapid, accurate and reliable recognition of chemical vapors is crucial for several applications in medicine[1-4], homeland security [5-8] and environmental monitoring [9]. To address challenges in chemical sensing, instruments combining an array of cross-selective chemical transducers with a pattern recognition engine, popularly referred to electronic noses or e-noses, have been proposed[10, 11]. Despite decades of efforts, these machine olfactory systems do not match the capability of their biological counterparts in terms of sensitivity and range of chemicals detected, as well as their stability over time [12, 13]. Several approaches to augment the performance of machine olfactory systems [14, 15], and even direct use of biological recognition elements (e.g. proteins, peptides) [16-20] or cultured cells/neurons[21-24] as transducers have been proposed. While these advances are significant, challenges remain in generating a rich repertoire of chemical transducers and translating the proposed approaches into low-cost, minimal-maintenance field-deployable units. This raises a following fundamental question: can e-noses with capabilities that match those of a relatively simple biological olfactory system, say an insect, be developed?

Even a relatively simple insect olfactory system employs on the order of 50 – 100 types of olfactory receptors [25]. There are several thousands of copies of each olfactory receptor neuron type thereby endowing the biological system with a diverse and large array of transducers. The biological sensory apparatus has been shown to detect diverse molecules, and some with exquisite sensitivity[26]. In addition, many design and computational principles in biological olfactory systems appear conserved in different organisms across phyla. One possible interpretation of this observation is that not many different solutions that are robust may actually exist. Further, considering that even a relatively small insect can possess a sophisticated chemical sensing apparatus compared to the state-of-the-art engineered e-noses, we sought to determine the feasibility of directly tapping into the capabilities of the insect olfactory system.

The envisioned approach is not that different from the ‘canary in a coal mine’ approach, where the viability of the entire organism is used as an indicator of absence/presence of toxic gases. Here, taking advantage of the modern electrophysiological tools, we demonstrate a simple stratagem. We let the biological transducers in the insect antenna convert the chemical signals into electrical neural signals which are then relayed to the olfactory centers in insect brain. We identified a bottle-neck region where the chemosensory information about volatile chemicals funnels in. Using a minimally-invasive surgical approach, we show that we can tap into the odor-evoked neural signals in this region in live insects. Multivariate responses across neurons in the recorded olfactory center are unique for an odorant, and therefore can be used as a fingerprint to recognize subsequent presentations of the same compound. As a proof-of-concept, we demonstrate that this part biological – part engineered system (i.e. a ‘biorobotic chemical sensing system’) can be used to rapidly detect and differentiate several different explosive vapors. Finally, we show how this concept can be extended to mobile robotic settings.

## Results

### Research hypothesis and the overall approach

In insects, volatiles chemicals (odorants) are transduced into electrical signals by olfactory receptor neurons (ORNs) present on the insect’s antenna (for example there are ∼50,000 ORNs in each locust antenna[27]). These odor-evoked electrical signals are then transmitted to the downstream antennal lobe (analogous to the vertebrate olfactory bulb). In the antennal lobe, the ORN input is reformatted by a circuitry comprising of two major types of neurons: ∼800 cholinergic projection neurons (excitatory; PNs) and ∼ 300 GABAergic local neurons (inhibitory; LNs)[28]. The PNs alone send their output wires, or axons, to two downstream centers: the mushroom body (∼50,000 Kenyon cells), and the lateral horn[28, 29]. As can be noted, the antennal lobe is a region where only the information about odorants funnel-in and are represented by a few neurons with higher signal-to-noise ratio[30, 31]. Therefore, the antennal lobe provides an ideal neural circuit to tap signals for acquiring information about the chemical cues present in the vicinity of an insect.

We chose the American locust (*Schistocerca americana*) for this study for several reasons:

i. they are sturdy and can recover from surgeries necessary for implanting electrodes
ii. their olfactory system has been very well studied[29, 31-33],
iii. they have non-spiking local neurons in the antennal lobe, therefore, signals from projection neurons alone can be monitored (PNs tend to be more odor specific than LNs),
iv. they can be trained to recognize odorants using classical conditioning assays[33, 34],
v. they can carry heavy pay loads, and,
vi. they can function in both solitary and gregarious phases (the latter can come in handy when multiple locusts or swarms are needed for remote sensing).

The overall approach taken in this study is schematically shown in **Fig. 1**. We hypothesized that locust antennal lobe neurons can be activated by diverse odorants, including many explosive chemicals and their precursors, that may not be ecologically important. Further, if responses from enough neurons are monitored simultaneously, useful stimulus-specific signal for detecting and recognizing diverse chemicals can be obtained (**Fig. 1A**). Integration with mobile robotic platforms would allow us to extend this idea to perform odor source localization tasks (**Fig. 1B**). In this study, we systematically investigated this issue, and developed experimental and analytical procedures to demonstrate the feasibility of this approach.

**Figure 1:**
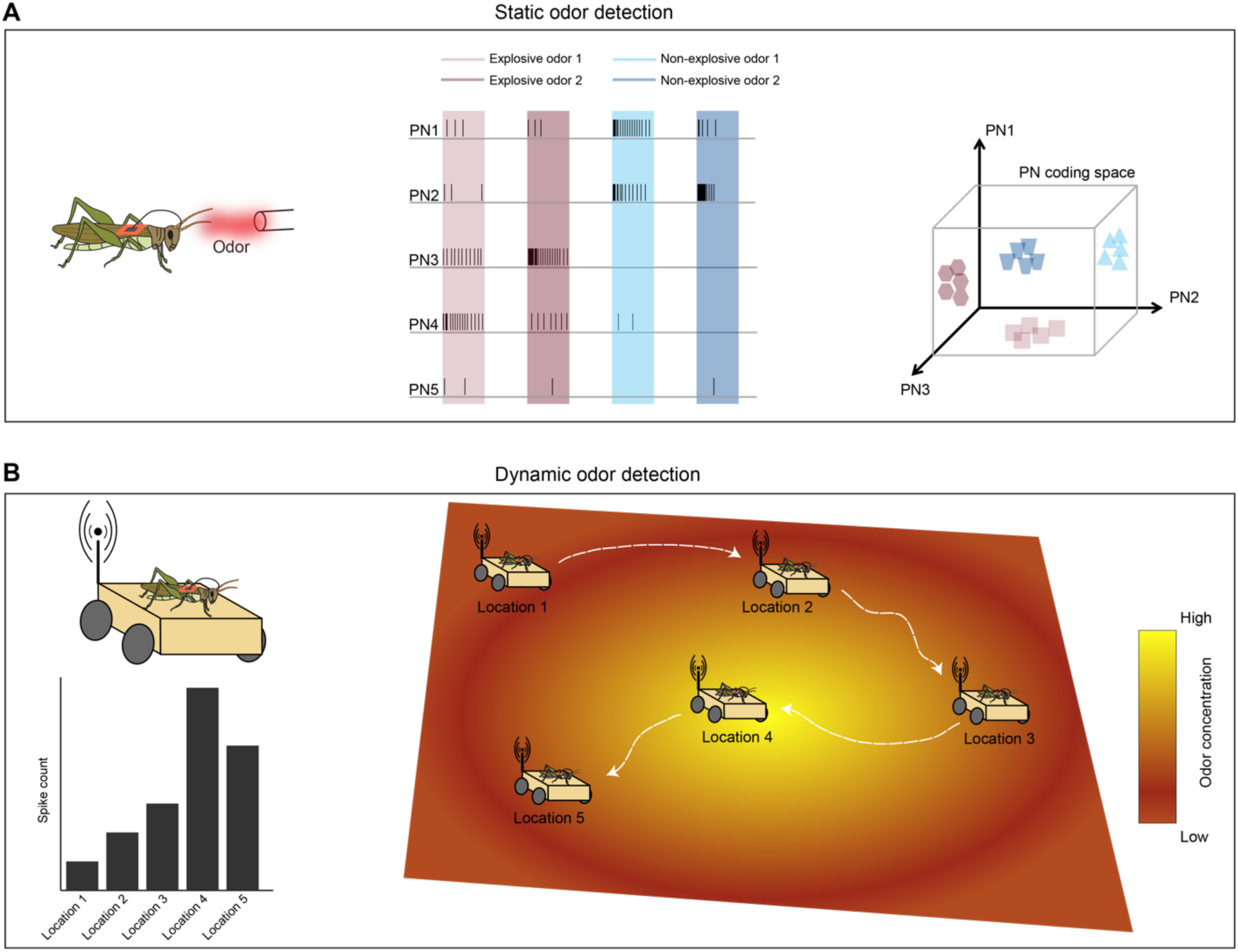
**A)** A schematic of the overall approach used in this study. The main idea is to hijack the insect olfactory system to perform chemical sensing. Left panel: Different odors are presented to the locust that has electrodes implanted in the olfactory regions of its brain. Recorded electrical activity is transmitted on to a backpack (shown in orange) that amplifies and records/transmits the signal. Middle panel: Schematic of how different olfactory odorants would elicit a stimulus specific response that can then be used to recognize the subsequent presentations of the same chemical. Note that each black line indicates one action potential, and the colored bars indicate odor presentation windows. In this schematic, projection neurons (PN) 1 and 2 are more responsive to non-explosive odors, PN3 and PN4 are more responsive to explosive odors, and PN5 is unresponsive to all odors. Right panel: Visualization of the representation of odor identity in a high-dimensional neural space. Note that responses elicited by different odors would occupy different regions as the ensemble responses across different neurons is unique and can encode for odor identity. **B)** An overview of using the insect olfactory system to decode odor identity in a dynamic setting. Left panel, top: A locust is implanted with electrodes in its olfactory circuits and the signal is recorded as in **panel A**. The locust is placed on a movable car and the electrical signal is transmitted wirelessly in real time. Right panel: Schematic of an open-field odor source localization task using the mobile locust and real-time monitoring of neural activity. Odor concentration in this region is shown as a heatmap with orange indicating low odor concentration and yellow indicating high concentration. The car is moved from location 1 to location 5 in a sequential fashion. Left panel, bottom: The spike counts recorded from the locust at each of the five locations. Note that high spiking activity would be expected near the odor source.

### Explosive odorants evoke discriminable responses in the insect brain

Do neurons in the locust antennal lobe respond to explosive chemicals that may be of potential interest to security applications? To determine this, we used a standard, highly-invasive surgical protocol[35], and implanted rigid, multi-unit, extracellular electrodes in the locust antennal lobe (**Fig. 2A**). As noted earlier, local neurons in the locust antennal lobe only fire calcium spikelets that are not detectable with the extracellular electrodes[27, 32]. We monitored the responses of individual projection neurons (PNs) to a panel of odorants, and each stimulus was puffed onto the antenna using a custom-designed olfactometer. The odor panel included hexanol (a green-leaf volatile of ecological importance), vapors of trinitrotoluene (TNT), its pre-cursor 2,4-dinitrotoluene (DNT), and hot air. Note that the DNT and TNT samples were purchased as solid crystals that required heating to create vapors (delivered concentrations in the tens to hundreds of parts-per-billion; see Methods). Since previous studies had shown that temperature alone can alter the spiking properties of the PNs [36], we included hot air as a control stimulus in these experiments.

**Figure 2:**
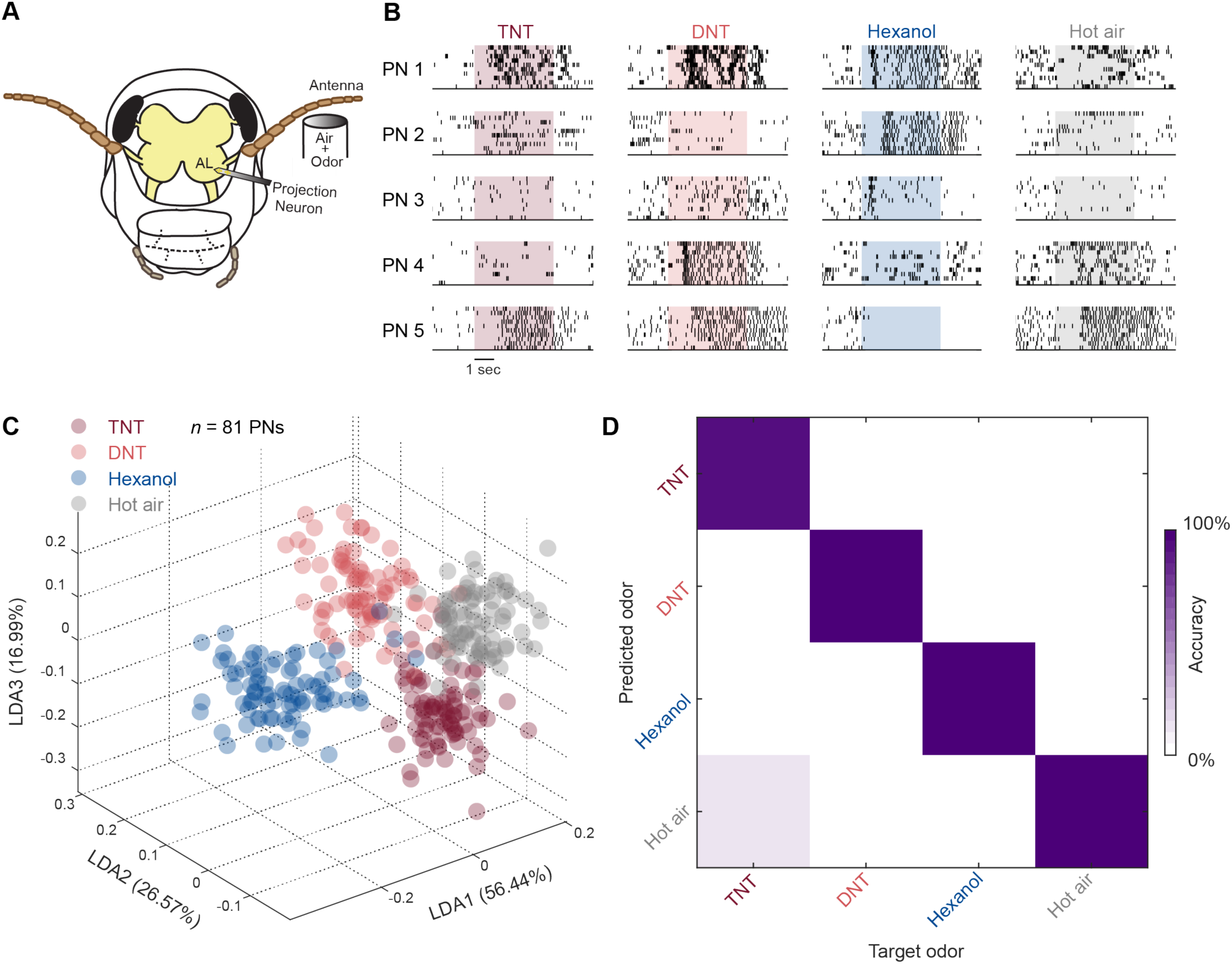
**A)** Schematic of the invasive surgical protocol for recording odor-evoked neural signals from the antennal lobe in the locust brain using rigid electrodes. Invasive protocol allows for precise access to the projection neurons (PNs) in the antennal lobe (AL). **B)** Raster plots of neural activity recorded from 5 projection neurons (PNs) in response to TNT, DNT, hexanol, and hot air (control) are shown. Each tick indicates a spike (or action potential) fired by the projection neuron and each row represents a trial. Shaded boxes indicate a 4 s time window when an odorant was presented to the antenna. Each column represents the responses evoked by a single volatile organic chemical across the same five projection neurons. Note that the five PNs respond differentially to different chemicals but the responses to any one stimulus is consistent across trials. **C)** Responses of 81 PNs during odor presentation window (50 ms binsize, over 4 s, total 80 points for each odor) are visualized after linear discriminant analysis (LDA) dimensionality reduction. Numbers within parentheses indicate the variance captured along that axis. The three dimensions onto which the data are projected, maximize the variance between classes and minimize within-class variance. The distinct clustering indicates the feasibility of segregating these chemicals based on the neural responses they elicited. **D)** Classification performance quantified via *leave one trial out* cross validation analysis is displayed as a confusion matrix. Each column corresponds to the target stimulus and row indicates predicted class. As most of the predicted responses match the target labels (diagonal elements), the information contained in the neural responses is sufficient to detect and recognize both explosive and non-explosive chemicals.

We found that some PNs responded to exposures of TNT and DNT vapors (**Fig. 2B**). Some responded to both these chemicals, while others responded preferentially to TNT or DNT. Hot air puffs also generated weak responses in some projection neurons, but these signals were different from those elicited by TNT or DNT. In many cases, the same PN responded to both the food odor (hexanol) and explosives (DNT and TNT), but with different firing patterns. These observations are consistent with the well-known broad response tuning properties of individual PNs in the locust antennal lobe[29, 32]. Notably, these results indicated that TNT and DNT activate subsets of olfactory receptor neurons in the locust antenna, just like other common chemicals that a locust would encounter (else there would be no detectable signals in the antennal lobe).

Is there enough information to reliably discriminate the two explosive chemicals from each other and from the other common odorants? It is well-established that in the locust antennal lobe, the odor identity is not encoded by single neurons. Rather, population neural responses and their evolution over time (i.e. response dynamics) uniquely represent individual chemicals[29, 32]. Therefore, we examined the information content at a population level. We combined the responses of all 81 projection neurons that were recorded to create an ensemble vector (see **Methods**). We then binned the activity into 50 millisecond time bins to provide a snapshot of the stimulus-evoked response across the entire population. Each odor exposure was 4 s in duration, so 80 trial-averaged snapshots of ensemble neural activity were obtained for each stimulus. To visualize the responses and compare them against one another, the high-dimensional responses were dimensionality reduced using linear discriminant analysis (**Fig. 2C**; LDA). As can be observed, each stimulus generated a unique neural response across the ensemble of PNs and therefore formed distinct response clusters after LDA dimensionality reduction. Note that TNT and DNT population neural responses were distinct from each other and were easily distinguishable from the responses evoked by the other stimuli. These dimensionality reduction results were supported by a quantitative classification analysis with leave-one-trial-out cross validation (**Fig. 2D**). Note that the confusion matrix is mostly diagonal indicating robust recognition of the different odorants. Overall, these results show that projection neurons in the locust antennal lobe respond to explosive chemical vapors and at an ensemble level carry information to support reliable recognition.

### Tapping neural responses in stable, mobile preparations

Having established the feasibility of the approach with a highly-invasive preparation (**Fig. 2**), we wondered if we could develop a general approach that would allow us to tap neural signals while insects explore complex environments. Further, we sought to optimize and establish a simpler surgical approach where the integrity of the organism is not compromised and most of its sensory and motor capabilities are retained. Such a preparation, we hypothesized, would allow long-term, stable recordings. Therefore, we developed a surgical procedure with minimal removal of the cuticle and tissue above the olfactory centers (**Fig. 3A**). We found that the locusts recovered from such minimally-invasive surgeries faster, and could move and feed without any hindrance. Further, we inserted a flexible, twisted-wire tetrode into the antennal lobe to monitor odor-evoked neural responses from the locusts which were no longer required to remain stationary. More importantly, neural responses recorded from such locusts implanted with electrodes were stable for long periods of time (see **Supplementary Fig. 1, 2**).

**Figure 3:**
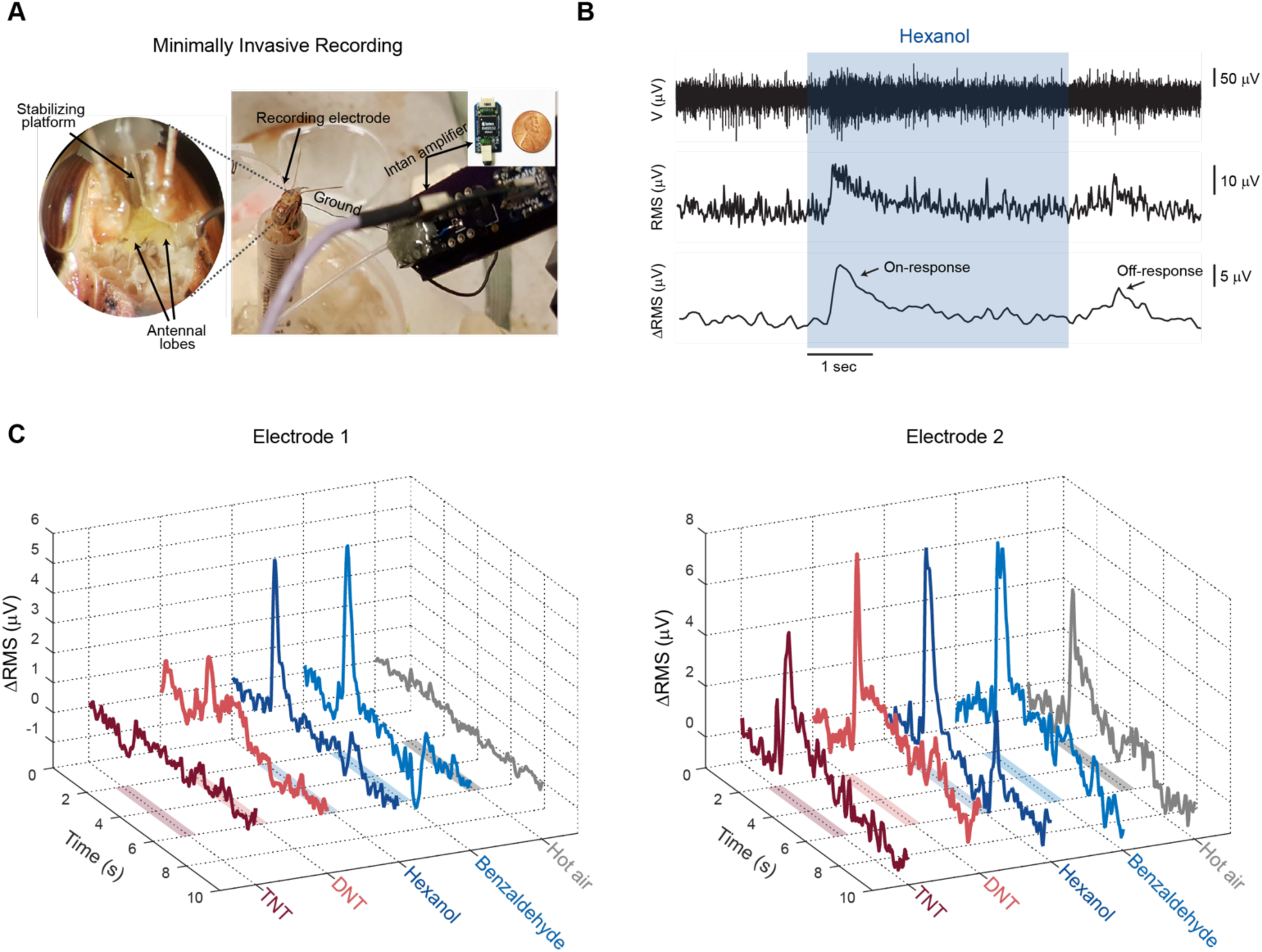
**A)** A picture of locusts with electrodes implanted into its antennal lobe using a minimally invasive surgery procedure is shown. The expanded panel shows the relative sizes of the implanted locust and amplifier (Intan Technologies). **B)** Calculation of signal energy from raw recorded neural activity. Colored rectangle indicates the time window when odor stimulus was presented. Briefly, it is obtained by calculating the RMS signal in a 50 ms moving window. The signals were smoothed and baseline subtracted. For details see **Methods.** **C)** Example traces of processed RMS signals recorded from two different electrodes are shown. Colored rectangles indicate time when odor was presented (4 s). RMS signal observed in different electrodes are distinct for different chemicals.

Note that each extracellular electrode monitored aggregated signals from many proximate neurons. Signals from multiple electrodes were used to separate the signal source (i.e spike sorting to assign each spike to an individual neuron). However, we noted that signals from several neurons were filtered out because of spike sorting as they were not clearly resolvable. Since these lost signals could also be potentially useful for target discrimination, we decided to use the total energy of the signal acquired from each electrode for detecting whether there is an odorant and determining its identity (**Fig. 3B**). This operation is both computationally cheap and ensures that all signals recorded are used towards discriminating the odorants. Further, total signal energy was monitored for each electrode in a 50 ms moving window, and averaged across trials. Measurements from different electrodes were concatenated to create a multivariate neural response vector that was used for evaluating the response specificity. **Fig. 3C** shows the unique RMS signatures evoked by different odorants in different recording electrodes.

We probed the responses of neurons in the antennal lobe as in the previous case while the locusts were exposed to two different odor panels (**Fig. 4A, Supplementary Fig. 4**). The first odor panel was similar to the one used in the previous set of experiments and included: TNT, DNT, hexanol, benzaldehyde and hot air. The second set had a larger and more complex set of odorants: RDX, ammonium nitrate, PETN, pATP, hexanol, benzaldehyde, acetonitrile, and hot air. **Fig. 4B, C** show dimensionality reduced neural response measurements made from locusts while they were exposed to various analytes in these two odor panels. Note that different chemical species evoked neural responses that were unique and different from the others indicating that every odorant used in the two odor panels, including the five different explosive chemicals, could be detected and precisely recognized (similar results for odor panel B shown in **Supplementary Fig. 4B, C**). Also, explosive and non-explosive chemicals forming distinct clusters when discriminability is considered for the broad categorical case (explosive vs non-explosive vs control; **Fig. 4B; also refer Supplementary Fig. 4B**). These qualitative dimensionality reduction analyses results were again verified using a quantitative classification analysis (**Fig. 4D, Supplementary Fig. 4D**). The confusion matrices were largely diagonal indicating low misclassification rates.

**Figure 4:**
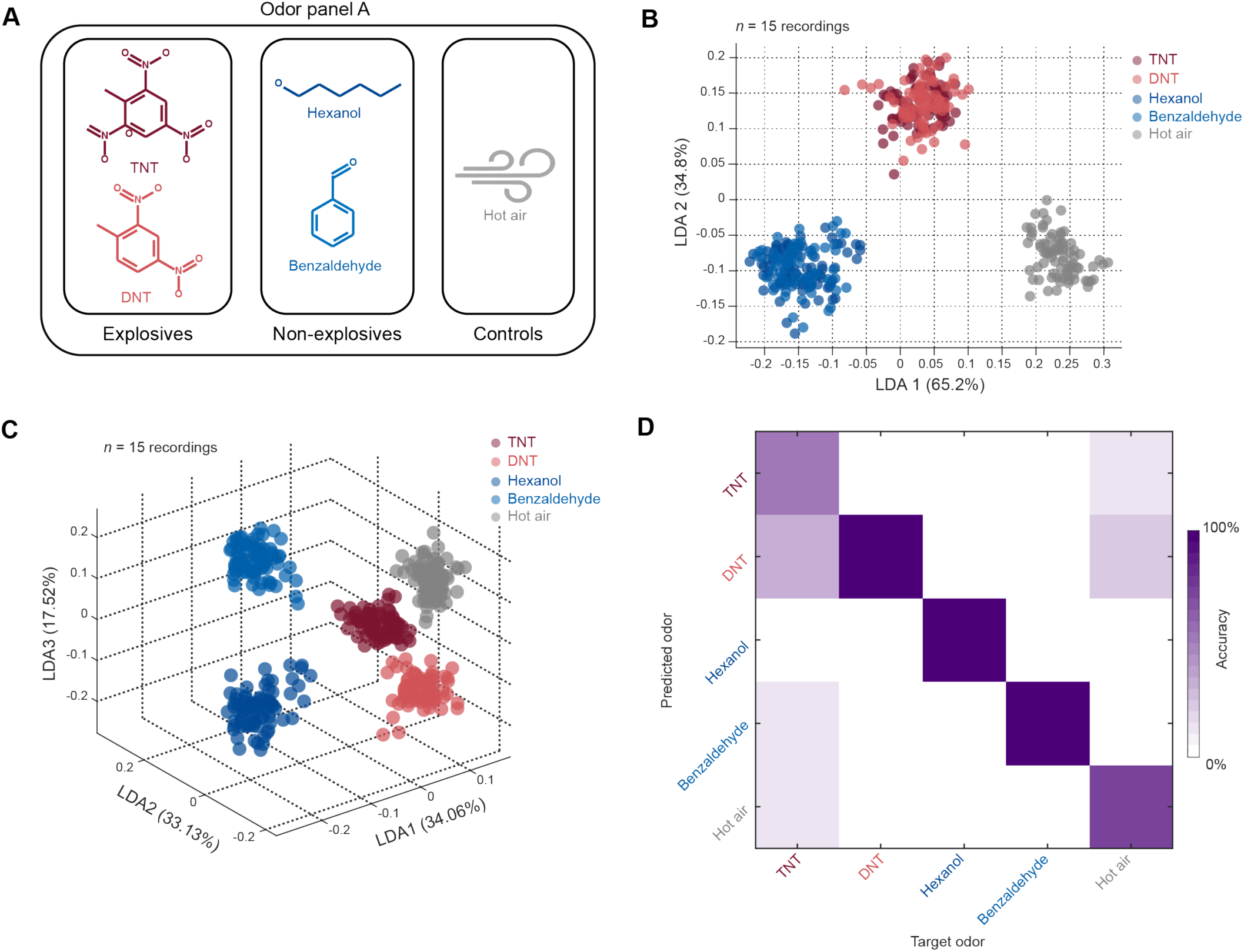
**A)** Two odor panels were used to probe neural responses in locusts implanted with electrodes using the minimally invasive procedure. The stimulus set comprised of explosive and non-explosive chemicals and appropriate control stimuli. Odor panel A included TNT, DNT, hexanol, benzaldehyde and hot air. TNT and DNT were in crystal form and were heated to 50 °C to generate enough vapors. Therefore, puffs of air heated to the same temperature was used as an additional control. **B)** Visualization of high dimensional data through dimensionality reduction using a 3-class linear discriminant analysis (LDA) is shown. Numbers within parentheses indicate the variance captured along that axis. Chemicals were labelled as explosives (red symbols), non-explosives (blue symbols) or controls (gray symbols). LDA shows clear separation between explosives and non-explosive chemicals (*n* = 15 recordings). **C)** Visualization of high dimensional data through a multi class LDA where each chemical was treated as its own class is shown. Numbers within parentheses indicate the variance captured along that axis. These plots show that individual explosives show distinct responses (*n* = 15 recordings for panel A). **D)** Confusion matrices summarizing the results from classification analyses are shown. Note that the matrices are mostly diagonal indicating that both explosive and non-explosive chemicals can be correctly identified based on the neural responses they evoke.

### Wisdom of the swarm

A major challenge in stand-off chemical sensing and localization is detecting target odorants which are heavily dispersed and subject to complex wind dynamics. Biological systems have evolved to mitigate these challenges. However, when tapping into these biological systems, there is a loss of information at each stage of the recording process. For example, we can record from only a subset of the neurons in the antennal lobe and increasing the number of recording sites could compromise the biological system. Data processing and classification lead to further loss in information. Using multiple locusts increases the likelihood of an ORN on the antenna getting activated by the target odorant and our recording system picking up the downstream activity in the antennal lobe. Therefore, we investigated whether increasing the number of locusts increases the signal-to-noise ratio and thus the classification accuracy of explosive chemicals. First, we performed classification analysis on single locusts, using signal from just one electrode (that had the highest variance for the duration of the experiment). We found that responses from any single locust had enough information to outperform a naïve classifier (**Fig. 5A**). However, the gains in classification accuracy came from the locust’s ability to robustly classify relevant naturally occurring odors for which they have highly tuned responses. Only one locust was able to classify with an accuracy greater than 60%. To quantify the performance of a population of locusts, we performed Monte Carlo simulations by selecting data from a random subset of locusts. Combining data from multiple organisms led to significant improvements in performance, with average accuracy reaching 80% with just 7 locusts (**Fig. 5B**). Thus, as can be expected, our results indicate that sensing with multiple organisms would lead to more efficient detection of the target chemical species.

**Figure 5:**
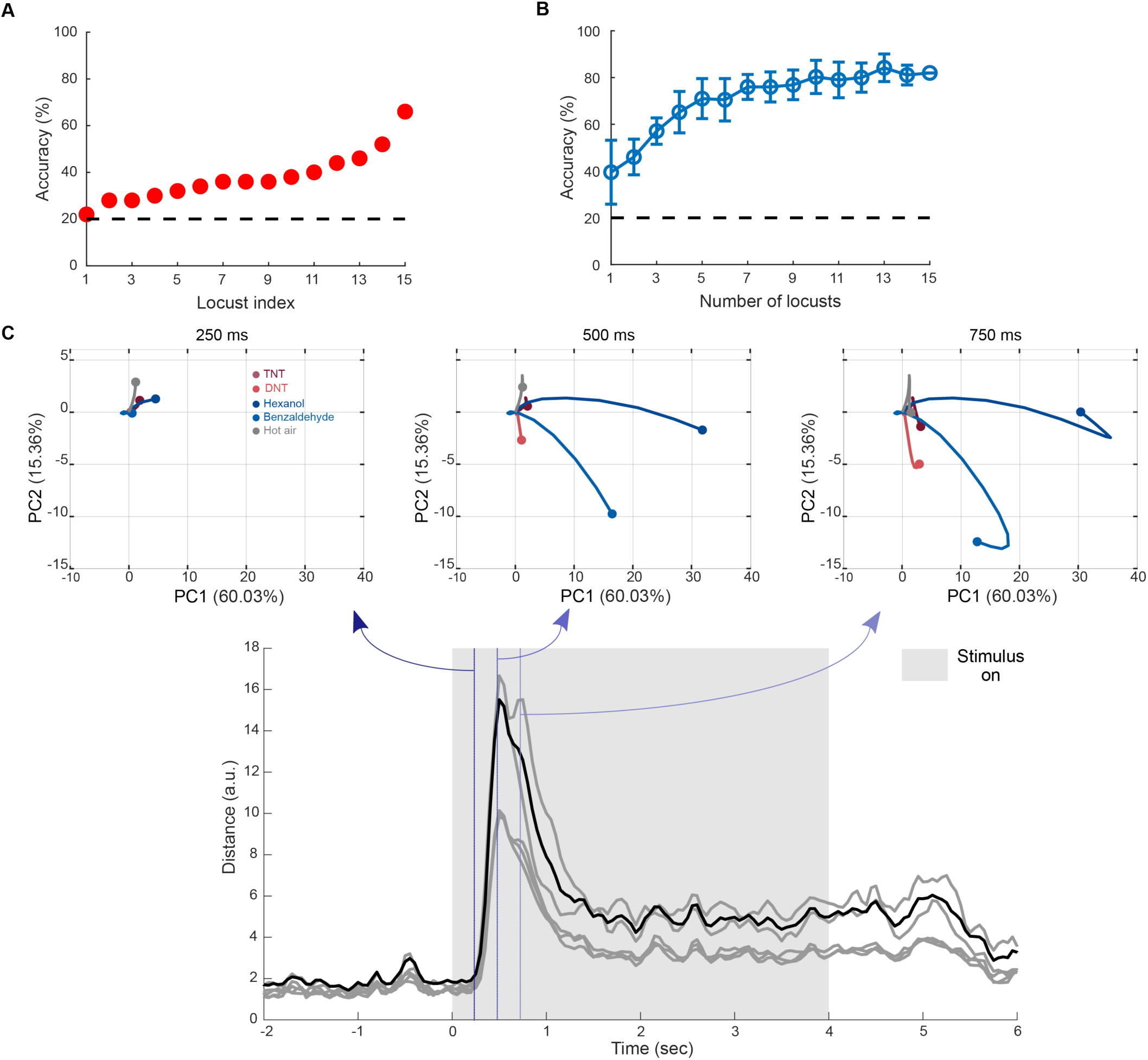
**A)** Classification performance using data recorded from the best channel/electrode in each individual locust is shown. Locusts are sorted based on the classification accuracy (low to high). Dashed line indicates performance of a naïve classifier for a 5-class problem (1 in 5 chance or 20%). **B)** Monte-Carlo simulations showing improvement in classification performance as data collected from multiple locusts are combined. Accuracy increases with number of locusts and reaches 80% using data from only seven locusts. **C)** Rapid identification of chemicals using neural responses: *Top panels*: high dimensional responses and how they evolve over time are visualized in two dimensions following principal component analysis. Numbers within parentheses indicate the variance captured along that axis. The three panels show evolution of neural responses for the first 250 ms, 500 ms and 750 ms after stimulus presentation. Note that neural responses are similar at 250 ms, but become distinct as they continue to evolve. Maximum separation is reached around 500 ms and and the responses start returning towards baseline within 750 ms. Thus, the transient neural responses can be utilized for rapid chemical identification. Bottom panel: pairwise distances between neural responses are plotted as a function of time. Black line indicates the mean pairwise distance across all chemicals. Maximum distance indicating maximum separation occurs at ∼500 ms after the onset of chemical stimuli.

### Rapid recognition of the targets chemicals

In many applications, rapid recognition of the target chemicals is highly desirable. Therefore, we sought to examine how quickly we could resolve the identity of the encountered chemical based on the neural signatures obtained. To understand how response patterns evolve over time, we performed a response trajectory analysis[32, 33]. For this analysis, the multivariate signal energy across electrodes were projected onto the top three eigenvectors of the covariance matrix (i.e. PCA dimensionality reduction; **Fig. 5C; Supplementary Fig. 3, 5**). Consistent with prior findings, both explosive and non-explosive odorants generated neural responses that evolved over time. The responses started from overlapping pre-stimulus baseline activity and quickly became odor-specific (**Fig. 5C**). We found that odorants became discriminable as the odor-evoked neural activity distributed across the ensemble of spiking neurons became odor-specific within 500 ms of their onset.

To verify this result, we computed pairwise distance between odorants and plotted them as a function of time (**Fig. 5C, Supplementary Fig. 5**). Consistent with the results from the PCA analysis, we found that the peak distance between pairs of odorants happened within 500 ms for all odor pairs (∼ 500 ms after stimulus onset for odor panel A and ∼350 ms for odor panel B). These results indicate that the neural response within a few hundred milliseconds of the odor onset is highly unique and can be used for rapid recognition of the chemical identity.

### Odor source localization

Finally, we examined if measurements made from the locust brain in controlled olfactory environments can be used to estimate concentration gradients; an important task for localizing the source of an odorant. For this purpose, we constructed an odor box where the concentration of a volatile chemical was varied along the length of the box (**Fig. 6B**). The odorant was introduced using an inlet port at the center of the box and a vacuum funnel directly opposite the inlet port removed the volatile chemical vapors. As a result, the odorant concentration was highest at the middle of the box and reduced exponentially on either side (**Fig. 6A**).

**Figure 6:**
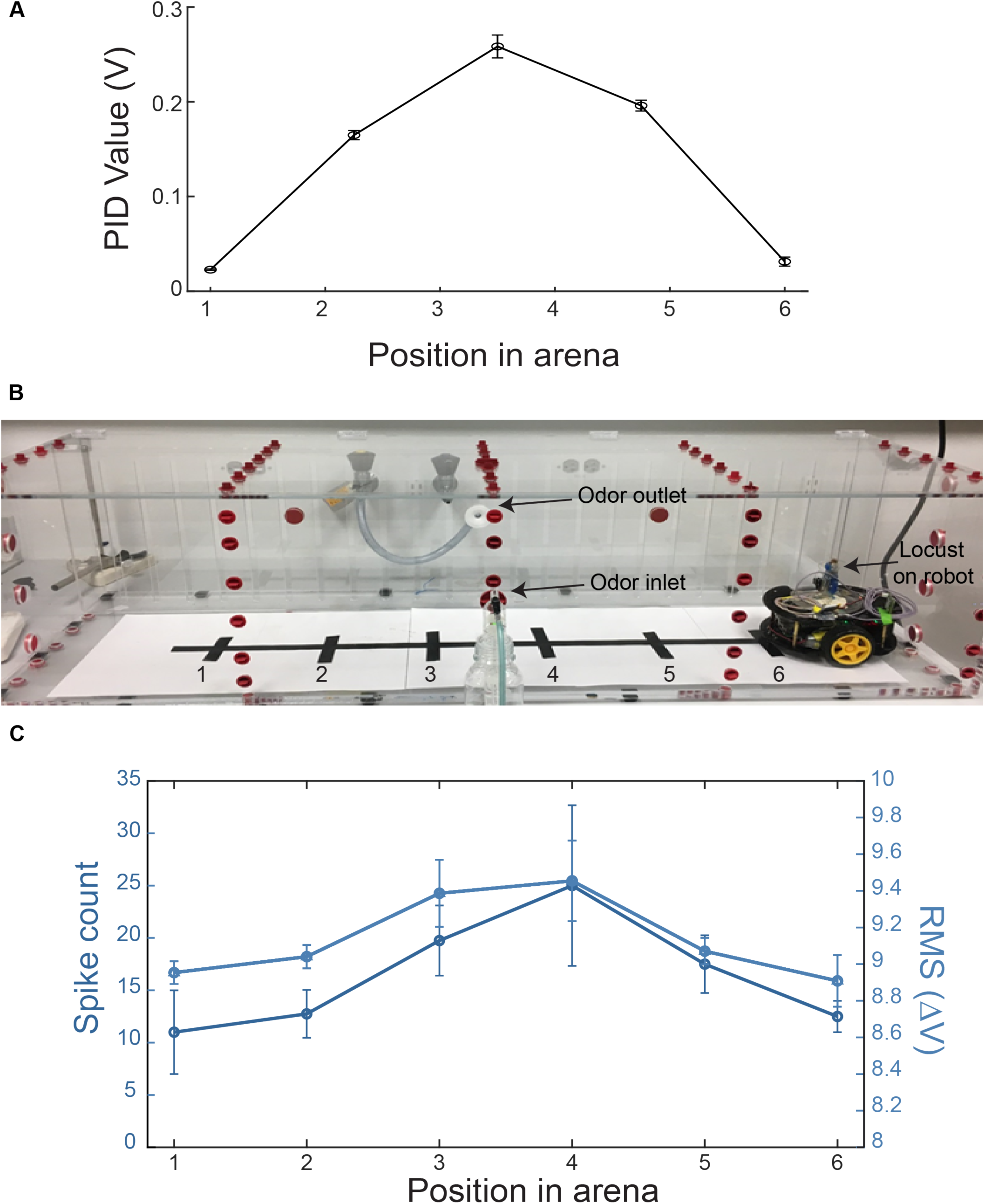
**A)** Quantification of odor gradient using a mini photoionization detector (mini200B, Aurora Scientific) inside a custom-made odor box designed to simulate an open-field environment. Odor concentration was found to be highest at the center of the box, with a symmetric drop on either side. **B)** Odor localization box designed for mobile experiments (see Supplementary Video 1). Odor is introduced into the box via the blue pipe near the front in the center, and is cleared out using a vacuum directly opposite to the inlet, shown in white. The car carrying the locust moves along the black tape running from right to left and stops at the 6 positions where tape runs across the main line. **C)** Spiking activity obtained from the locust is shown as a function of position in the arena are shown (see **Methods**). Spiking activity was recorded for periods when the car was allowed to remain stationary at the 6 positions for 20 s each. Mean spiking activity (left axis) and RMS signal energy (right axis) are shown across different runs, and the error bars indicate s.e.m. As can be seen, the locust spiking activity is highest in the center near the odor inlet (source) and drops off symmetrically, similar to the concentration profile of the odorant shown in **panel A**. (*n* = 2 for positions 1, 6; *n* = 4 for positions 2-5).

To estimate concentration, we implanted the electrodes into locust antennal lobe using the minimally invasive surgical protocol. We placed the implanted locust and associated neural recording amplifiers and a wireless transmitter in a mobile robot to monitor spiking activity changes as the locust was moved about in the odor box. Since we found that there were considerable motion artefacts in the recorded signals while the robot was moving, samples were acquired in a stop-collect-move fashion, as the mobile robot was stopped at fixed intervals along the major axis of the odor box (see **Supplementary Video 1**). Our results show that the spiking activity was greatest at the center of box and tapered as the robot moved towards corners on either side (**Fig. 6C**). Hence, the spiking rates varied with and matched the concentration profiles.

Taken together, these results provide proof-of-concept data to show that the neural signals from the insect brain could be tapped to resolve the identity of odorants in their vicinity and thereby can be used to perform remote chemical sensing.

## Discussion

We have demonstrated an approach where the sensing capabilities of a whole organism can be tapped to achieve a hybrid chemical sensing system. Transduction of chemical vapors into electrical signals were still achieved using the sophisticated biological olfactory receptor neurons (ORNs) in the insect antenna. Since each of these ORNs is noisy and there are several receptor neurons with diverse tuning (∼50,000 in each antenna), directly tapping their responses would not provide a robust approach. However, the outputs of ORNs conveniently funnel-in their information onto ∼800 downstream neurons in the antennal lobe. Each neuron in the antennal lobe responds to many odorants and with high signal-to-noise ratio. Therefore, we tapped into this antennal lobe region to record neural signals from multiple neurons simultaneously and use this for odor recognition. Our results reveal that indeed this approach is feasible and can provide rapid and reliable discrimination between vapors of different explosive chemical species.

For the goals of this study, we required the insect species chosen to be sturdy and with easy access to their olfactory regions in the brain. Amongst many insect species, *Periplaneta americana* (American cockroach) and *Schistocerca americana* (American locust) emerged as the viable candidates that satisfy these requirements. In locusts, only the projection neurons (PNs) fire full blown sodium action potentials, whereas local neurons fire calcium spikelets that are not detected with extracellular electrodes. It is worth noting that local neuron responses carry less information for odor discrimination. Given that the locust olfactory system has been well-studied, and only PN responses that carry rich odor discriminatory information can be tapped easily in this model organism, we chose this invertebrate model for achieving the goals of our study.

Prior works have demonstrated that neural signals such as electroantennogram, which monitors the total activity of the all ORNs[37, 38], or behavioral read-outs such as proboscis extension reflex[39], can be used for achieving chemical sensing with invertebrates. We note that the electroantennagram signals, while experimentally easy to acquire, tend to be less discriminatory overall. When a larger panel of chemicals need to be discriminated, electroantennogram may not provide discriminatory information to allow robust recognition. Further, we have noted that even odorants that do not evoke a strong EAG responses, elicit distinct and dynamic patterns of neural activity in the antennal lobe. Therefore, directly tapping neural signals from the antennal lobe may provide a better approach for realizing a hybrid chemical sensing system. With regards to behavioral readouts, we note that factors such as cross-generalization, presence of other confounding non-chemosensory stimuli, and variance in capabilities/performance across different individual insects may diminish the overall capability achieved. Also, some odorants may not be compatible with appetitive conditioning assays typically used to train these insects, thereby potentially limiting the utility of such readouts.

While our results reveal feasibility in tapping neural signals from the insect brain, several challenges remain to be overcome. First, we found that the overall recording quality remained intact for ∼7 hours (**Supp. Fig. 2**). The spike rates reduced for even longer recordings on the order of several days (**Supp Fig. 1**). We note that while the stability can be potentially improved by feeding locusts at regular time intervals, long-term operations beyond a few days may require further refinements to surgical techniques and materials used as electrodes that interface with the neural tissue.

The second issue concerns the transfer of training data obtained from one locust to another. Since the electrodes are placed randomly within the antennal lobe, it would not be possible to use data collected from one locust to perform odor recognition using signals obtained from a different locust. However, since the responses patterns are stable over a recording session, it would be possible to acquire the training data for target chemicals at the beginning of each recording session. Also, the easy availability of the insects for experimental manipulation, their economy and the practicality of the approach, we believe, offsets the other drawbacks.

## Acknowledgements

We thank members of the Raman Lab (Washington University in St. Louis) for feedback on the manuscript. Sarah Widder and Christian Pederson are acknowledged for their initial efforts in developing a minimally invasive recording preparation. This research was supported by Office of Naval Research grants (N00014-16-1-2426, N00014-19-1-2049) to B.R, S.C and S.S.

## Author contributions

BR conceived the study and designed the experiments/analyses. EA demonstrated initial feasibility using the invasive preparation. DS and RC independently developed minimally invasive recording approaches. DS showed feasibility of the overall approach and initial mobile robotic sensing experiments. DM setup the acquisition system. RC performed stability analysis, repeated experiments and validated the results. DM performed all the analysis. DS, DM and RC generated all the figures and co-wrote the methods section. MT developed the odor box and RL programmed the line-following robot. PG calculated the concentrations of the explosive vapors delivered in our experiments. SS provided the explosive chemical samples. SC advised on the instrumentation aspects of the work. BR wrote the paper taking inputs from all the authors and supervised all aspects of the work.

## Methods

### Surgical procedure

Post-fifth instar locusts (*Schistocerca americana*) of either sex raised in a crowded colony were used for all experiments. Locusts were immobilized with both antennae intact. The olfactory regions of their brain were exposed, desheathed following treatment with protease, and superfused with locust saline at room temperature. Extracellular, multiunit recordings of projection neurons (PN) were performed with a 16-channel, 4×4 silicon probe (NeuroNexus) superficially placed on the antennal lobe. These neural recordings procedures have been used in previous studies, and a visual demonstration of each of these steps is available online [35].

In addition to the standard procedure[32, 33, 35, 40], we developed a novel technique to perform neural recordings while minimally damaging the neural tissue and other organs (e.g. keeping antennae, muscles, maxillary palps, legs, and wings intact). In this minimally-invasive surgical technique, only a minor incision in the head cuticle was made to expose the antennal lobe of the brain (**Fig. 3A**). This small incision ensured that the locusts could still move their mouthparts and antenna freely. The brain was stabilized with a micro-platform not hindering mouth and palp movements and the antennal lobe was de-sheathed without the use of any enzyme. A custom-made, flexible, multi-wire electrode array that was inserted into the antennal lobe through the incision in the cuticle. A silver-chloride ground wire was also inserted into the head and secured with batik wax. A micromanipulator was used for the precise placement of the electrode while voltage signals were constantly monitored for optimal placement. Once positioned, the micromanipulator was removed and the electrode-array was held in position using batik wax. Neural signals were amplified and digitized using a miniaturized amplifier (Intan Recording System, RHD2132 16-Ch headstage). The entire setup was placed inside a custom-made Faraday cage and placed on a vibration isolation table or on a mobile robotic platform.

### Odor panel

For the invasive recordings, we delivered olfactory stimulation using a standard procedure [32, 33, 35, 40]. Briefly, odorants were first diluted in mineral oil to either 1% concentration by volume (v/v) and sealed in glass bottles (60 ml) with inlet and outlet ports. A constant volume (0.1 L/min) of the static headspace above the diluted odor-mineral oil mixture was displaced into a desiccated carrier air stream (0.75 L/min) using a pneumatic picopump (WPI Inc., PV-820). For all experiments, other the mobile robotic ones, a vacuum funnel placed behind the locust preparation continuously removed the delivered odors. Each odorant was presented for 4 seconds and was repeated ten times with 60 seconds inter-trial interval.

We used three odor panels for experiments reported in this study:

1. Invasive recordings: TNT, DNT, hexanol and hot air
2. Minimally-invasive recordings
  a. Odor panel A: TNT, DNT, hexanol, benzaldehyde and hot air
  b. Odor panel B: TNT, DNT, PETN, pATP, hexanol, benzaldehyde, acetonitrile and hot air

Solid TNT and DNT crystals were heated to 50 °C to generate vapors. RDX and PETN were dissolved in acetonitrile.

The concentration of the chemicals delivered to the locust was estimated by first calculating the vapor pressure of the chemical in the odor delivery bottle. Vapor pressure was calculated in the following ways:

1. For solid chemicals at room temperature (Ammonium nitrate, pATP) vapor pressure was obtained from [41] (also refer https://www.lookchem.com/4-Aminothiophenol/)
2. For heated solid chemicals (TNT, DNT), vapor pressure was obtained using Clausius–Clapeyron equation and obtaining the relevant parameters from [41]
3. For chemicals dissolved in a solvent (hexanol, benzaldehyde, RDX, PETN), the mole fraction of the chemical was calculated (https://pubchemdocs.ncbi.nlm.nih.gov). Partial vapor pressure was estimated by multiplying its vapor pressure by its mole fraction (Raoult’s law). Note that for RDX and PETN, due to their low vapor pressure, only rough estimates could be made.

Note that the concentration of the chemical in the bottle was obtained by normalizing its pressure with the atmospheric pressure. As this mixture was introduced @0.1 l/min into a continuous air stream of 0.75 l/min, the final concentration, incident on the locust antenna, was obtained by multiplying the bottle concentration by a factor of 1/8.5.

| Chemical | CAS Number | Delivery Method | Concentration<br>(in parts per million or parts per billion) |
| --- | --- | --- | --- |
| TNT | 118-96-7 | Solid, heated to 50 °C | 30.6 ppb |
| DNT | 121-14-2 | Solid, heated to 50 °C | 779 ppb |
| Ammonium Nitrate | 6484-52-2 | Solid crystals | 15.1 ppb |
| RDX | 121-82-4 | 1mg/ml dissolved in acetonitrile | < 1 ppb |
| PETN | 78-11-5 | 1mg/ml dissolved in acetonitrile | < 1 ppb |
| Hexanol | 111-27-3 | 1% v/v dissolved in mineral oil | 4.44 ppm |
| Benzaldehyde | 100-52-7 | 1% v/v dissolved in mineral oil | 7.46 ppm |
| pATP | 1193-02-8 | Solid powder | 1.7 ppm |
| Acetonitrile | 75-05-8 | Liquid solvent - control | 0.013 |
| Air |  | Heated to 50 °C for panel A, room temperature for panel B | 1 |

### Data collection and processing

Data collection and analyses for the three sets of experiments – invasive preparation and odor panels A, B for minimally invasive preparation – were done separately. For the invasive prep, extracellular neural data were collected using a NeuroNexus 16-channel, 4 × 4 silicon probe. Impedances for each electrode contact pad was maintained in the 200–300 kΩ range via gold electroplating prior to each experiment. Signals were filtered between 0.3 and 6 kHz and amplified by a 10k custom-made 16-channel amplifier (Biology Electronics Shop; Caltech, Pasadena, CA). They were then digitized at 15 kHz using custom LabView Software. For each odor, 10 repeated trials were performed. The collected signals were spike sorted as described previously [33, 40, 42]. We identified 81 PNs from recordings obtained from 11 locusts (16 antennal lobes). Spikes were counted and binned in 50 ms time-bins.

For minimally invasive surgery, we used custom-built 8-channel twisted wire electrodes. We observed that not all channels were electrically active. We collected data only from those electrodes that had impedances impedance between 100kΩ and 500 kΩ and the sampling rate was 15 kHz. In total, we collected data from 65 channels (across 15 locusts) for odor panel A and 33 channels (across 6 locusts) for odor panel B. 10 repeat trials were performed for panel A, while 5 trials were carried out for panel B.

### Signal energy calculations

We observed that spike sorting led to heavy loss of information and used an alternative data processing approach (signal energy) for pre-processing signals collected using the minimally-invasive procedure. The collected raw data were filtered using a bandpass filter between 300-5000 Hz and passed through a continuous moving RMS filter with a 20 ms window size (using standard Matlab DSP toolbox). The data were then down-sampled by a factor of 150, smoothed by a ten-point moving average filter and further down-sampled by a factor of 5. The final temporal resolution that was used for rest of the analysis was 50 ms (same temporal resolution as the invasive preparation).

We defined the baseline RMS signal level for each trial by taking a mean of the RMS signals observed in a two-second pre-stimulus window. To obtain the odor-evoked responses, baseline RMS voltage was subtracted to obtain the ΔRMS values (**Fig. 3B**).

### Dimensionality reduction analyses

For data visualization, we performed two kinds of dimensionality reduction analyses – Linear Discriminant Analysis (LDA) and Principal Component Analysis (PCA). In both cases, only responses during presence of stimulus (4 sec; i.e. 80 time bins) were considered. For all analyses, we constructed a response matrix in the following fashion. First, responses collected from different locusts were aligned with respect to the odor onset and averaged across trials. This resulted in a block of 80 columns that represented the trial-averaged responses observed in different locusts/electrode combinations (each row) to a single odorant. Responses to different odorants were concatenated. For example, if there were data from five different locusts (one electrode each) to 4 different odorants analyzed, the resulting data matrix would be: 5 rows times 320 columns (4 × 80). For the three experiments we performed, we ended with the following data matrix dimensions:

- Proof-of-concept panel (**Fig. 2**): 81 PNs x 320 time bins (4 odors),
- Odor panel A (**Fig. 4**): 65 channels x 400 time bins (5 odors)
- Odor panel B (**Supplementary Fig. 4**): 33 channels x 640 time bins (Panel B – 8 odors).

Note that each column corresponded to a point in *d*-dimensional space, with *d* being number of total channels (or number of PNs) recorded for each panel of odors.

For multi class LDA, each column vector was labeled based on the stimulus identity, while for 3 class LDA, it was labelled as the group to which the odor belonged – explosives, natural volatiles and controls. Within class variance S^(w)^, and between class variance S^(b)^, were estimated.

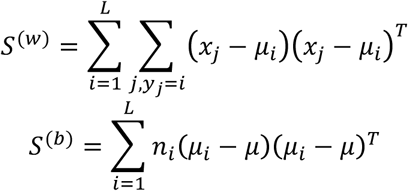

Where x_i_ is *i*^th^ column vector in data matrix, y_i_ is the label of that point, L is number of classes, μ_i_ is mean of points belonging to class *i*, μ is mean of all points and *n*_*i*_ is number of points belonging to class *i*. Eigenvectors and corresponding eigenvalues of the matrix (S^(w)^)^-1^*S^(b)^ were calculated. The data points were then projected onto two or three eigenvectors corresponding to the largest eigenvalues (two eigenvectors in case of LDA with 3 classes, three for multi class LDA). These eigenvectors maximize distance between classes, while minimizing within-class variance. The projected points were color-coded by their stimulus (**Fig. 2C, 4B, C, Supplementary Fig. 4**).

Principal Component Analysis (PCA) was used to visualize how ensemble neural responses change over time (i.e. a trajectory in state space). For PCA dimensionality reduction, we calculated the data covariance matrix and projected the data onto two (**Fig. 5C, Supplementary Fig. 5**) or three (**Supp. Fig. 3**) principal eigenvectors that captured maximum variance in the data. During plotting, the odor trajectories were created by joining temporally consecutive points after smoothing using a five-point moving-average, low-pass filter.

### Classification analysis

Observations were labelled as described for the LDA analyses (see above). For obtaining an unbiased estimate of classification accuracy, we performed *leave one trial out* cross validation analysis. During each iteration, data from one trial for all odors were removed from the training set and used as test data. A quadratic discriminant was fit on the remaining data where the predictors were the neural responses in one time-bin and the expected classifier output was the class label for that time-bin. Thus, for a test trial for one odor, we obtained 80 predicted responses. The class label for a trial was taken to be the mode of those 80 responses. A confusion matrix was created by comparing the predicted responses to the known responses. Briefly, *C*_*ij*_ is the number of trials of Odor *i*, predicted to be Odor *j*. A fully diagonal matrix indicates 100% classification accuracy.

### Locust population analysis

For determining how performance varied as a function of number of locusts used in the analyses (**Fig. 5B**), a random subset of locusts were chosen (from n locusts choose k random locusts; k was varied) and signals from one electrode with the maximum variance was selected for each locust. The signals from different locusts were combined and the classification analyses was repeated. Accuracy was calculated as ratio of correctly classified trials to total trials. This was repeated 20 times, and the mean and standard deviations of accuracy for each group size were calculated and plotted in **Fig. 5B**.

### Recordings on a mobile robotic platform

After implanting electrodes into the locust brain, we sealed-off the incision made to expose the brain, and moved the implanted locust preparation to a mobile robotic platform. The locust was secured inside a plastic tube, which also worked as a harness for the electrode base, micro-amplifier and wireless transmitter (i.e. back-pack). This preparation was moved back and forth in the odor box to sample chemical vapors (**Fig. 6B**). The locust was fully exposed to its environment since no Faraday cage was used. Neural signals were digitized and amplified at the backpack attached to the locust. The back-pack was connected to the computer via a wireless transmitter (RCB-W24A-LVDS, DSPW) which gave the preparation total mobility.

### Characterizing chemical gradient in the odor box

We used a fast photo-ionization detector (mini200B, Aurora Scientific) to characterize odor distribution in the odor box. The PID was placed at uniformly spaced locations within the box and data were acquired for five repeated presentations of the odor at each location. Raw data were acquired at 15 kHz sampling rate using a custom MATLAB program. The mean of the peak voltage signal across trials was used to represent the odor concentration at each sampled location. The mean signal and the s.e.m across trials is shown in **Fig. 6A**.

### Locust responses in the odor box

We used a simple threshold-based event detection approach to count the number of supra-threshold events, where each event was a putative spike. A threshold of 8 s.d. above the baseline was used to detect the events. Using this approach, we counted the number of events or spikes at each location and the mean spiking activity at each stationary location along with the s.e.m. is plotted in **Fig. 6C**.

---

**Supplementary Figure 1:**
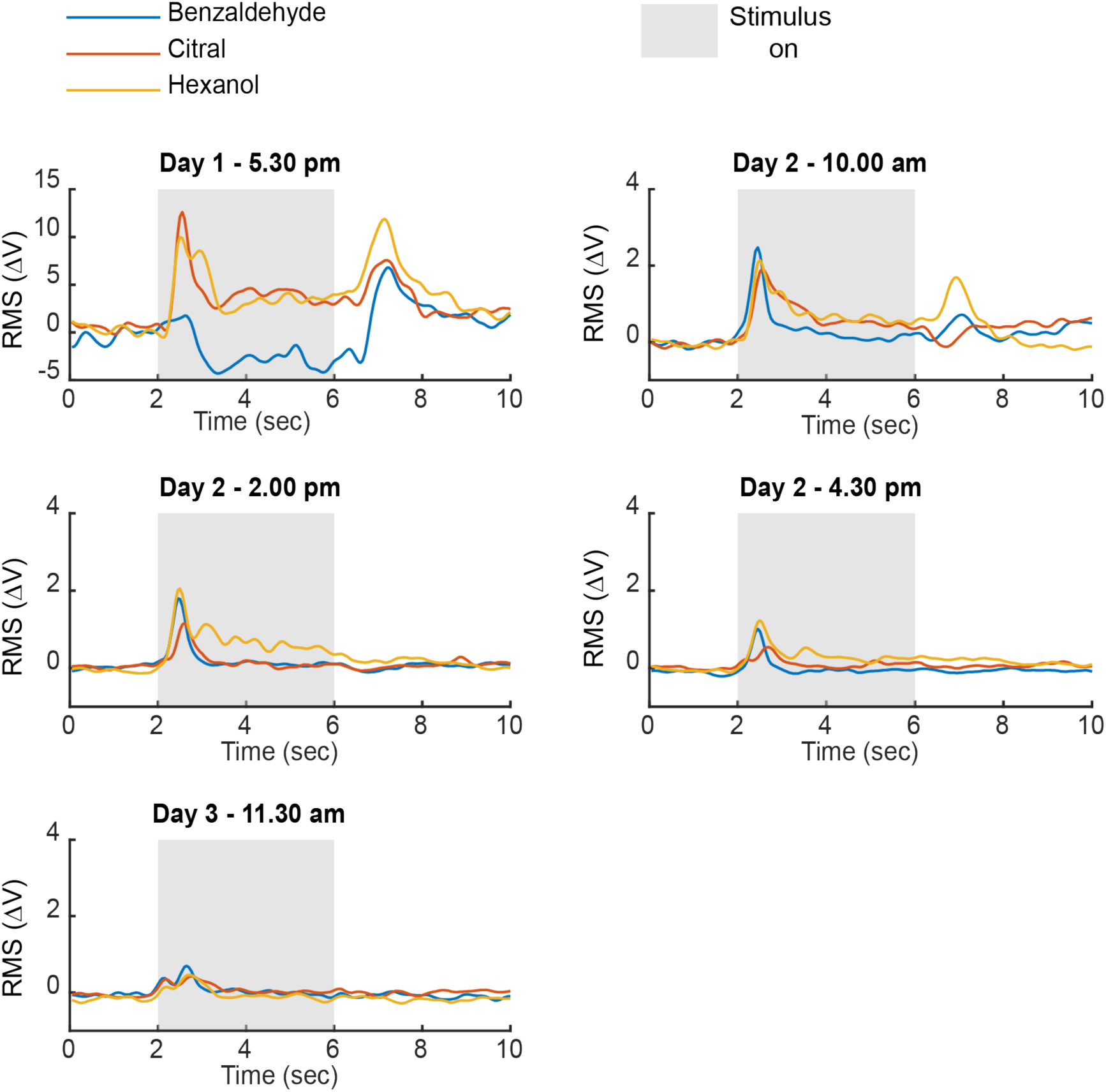
Long-term recording using the minimally invasive surgical protocol. Neural responses (RMS signal energy) to three odorants: benzaldehyde, citral, and hexanol are shown. Grey rectangle indicates the 4 second period when the stimulus was presented. An increase in the signal energy at stimulus onset can be seen consistently from Day 1 (5:30pm) to Day 3 (11.30am), a period of 42 hours.

**Supplementary Figure 2:**
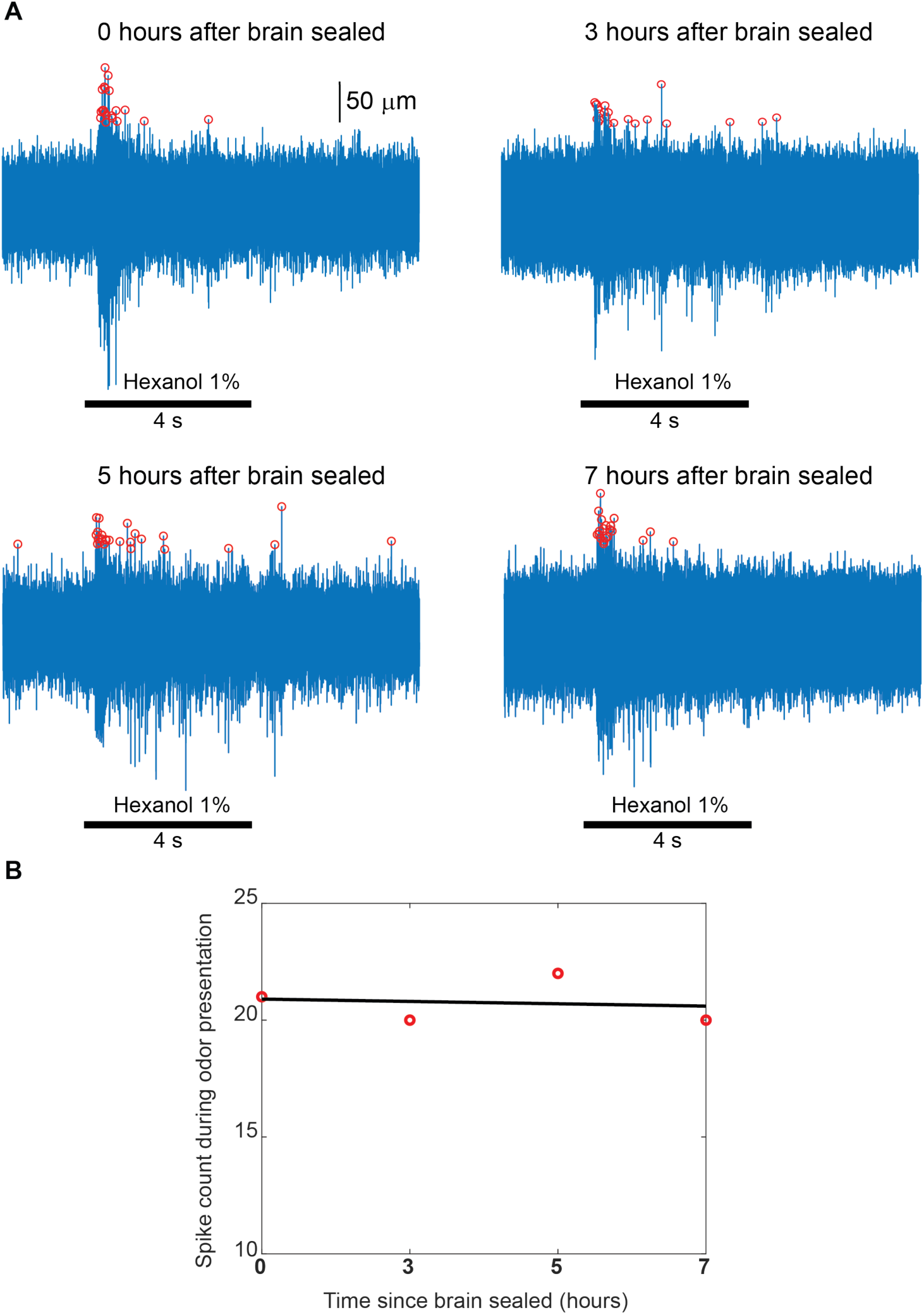
**A)** Long-term recording using the minimally invasive surgical protocol. Neural responses to presentations of hexanol (odorant) are shown as raw extracellular signals. Spikes identified using a threshold detection approach (see **Methods**) are indicated using red circles. Robust and consistent spiking responses to the presentation of the odor can be seen for up to 7 hours after the brain was sealed post electrode implantation. **B)** Spike counts (red circles) during the 4 seconds of odor presentation are shown as a function of time. The best-linear fit is shown in black. Note that the odor-evoked responses remain consistent for the entire duration of the experiment.

**Supplementary Figure 3:**
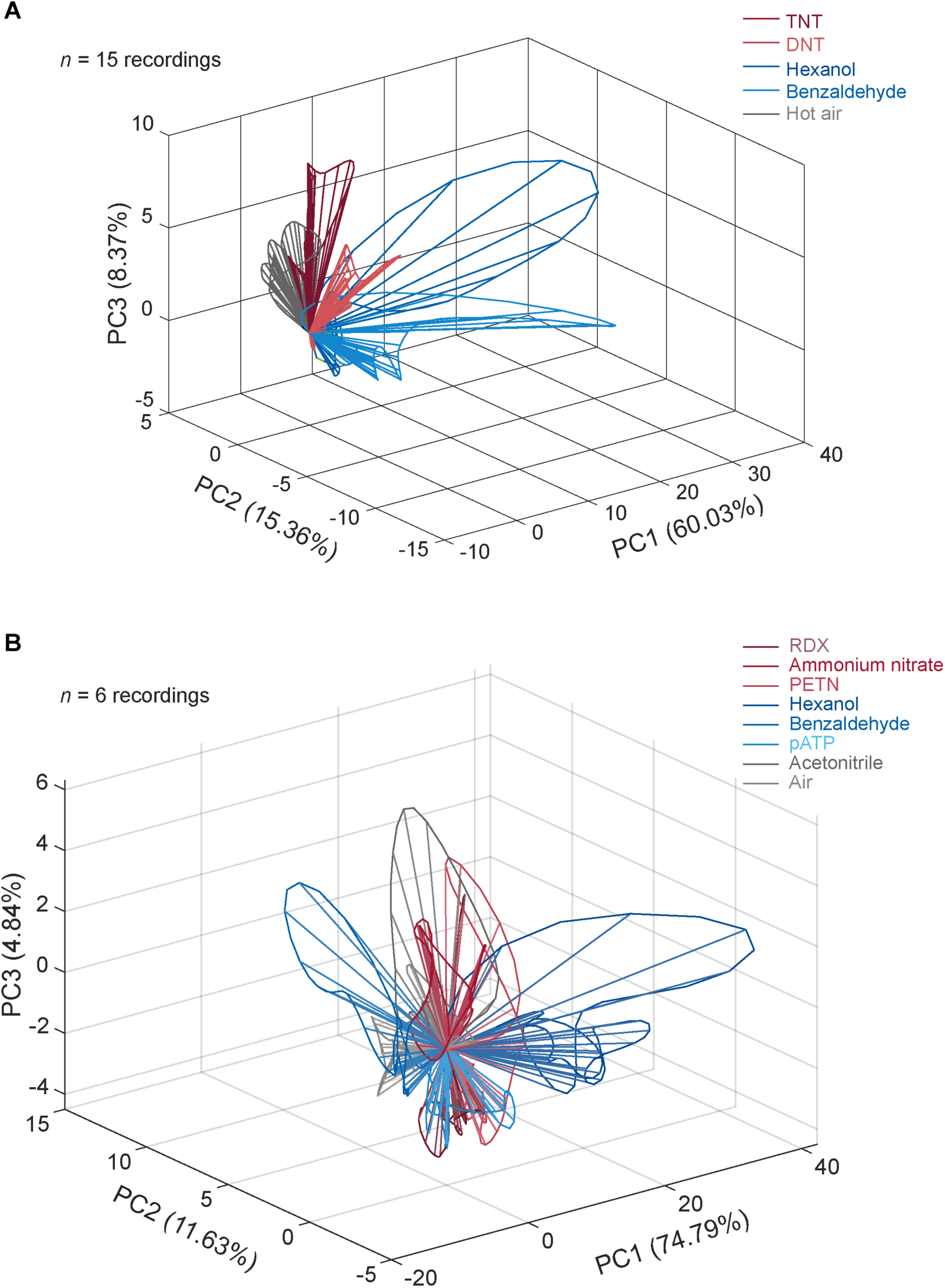
**A)** High dimensional responses for odor panel A and how these responses evolve over time are visualized in three dimensions following principal component analysis (see **Methods**). Numbers within parentheses indicate the variance captured along that axis. **B)** Similar analysis but showing results for odor panel B.

**Supplementary Figure 4:**
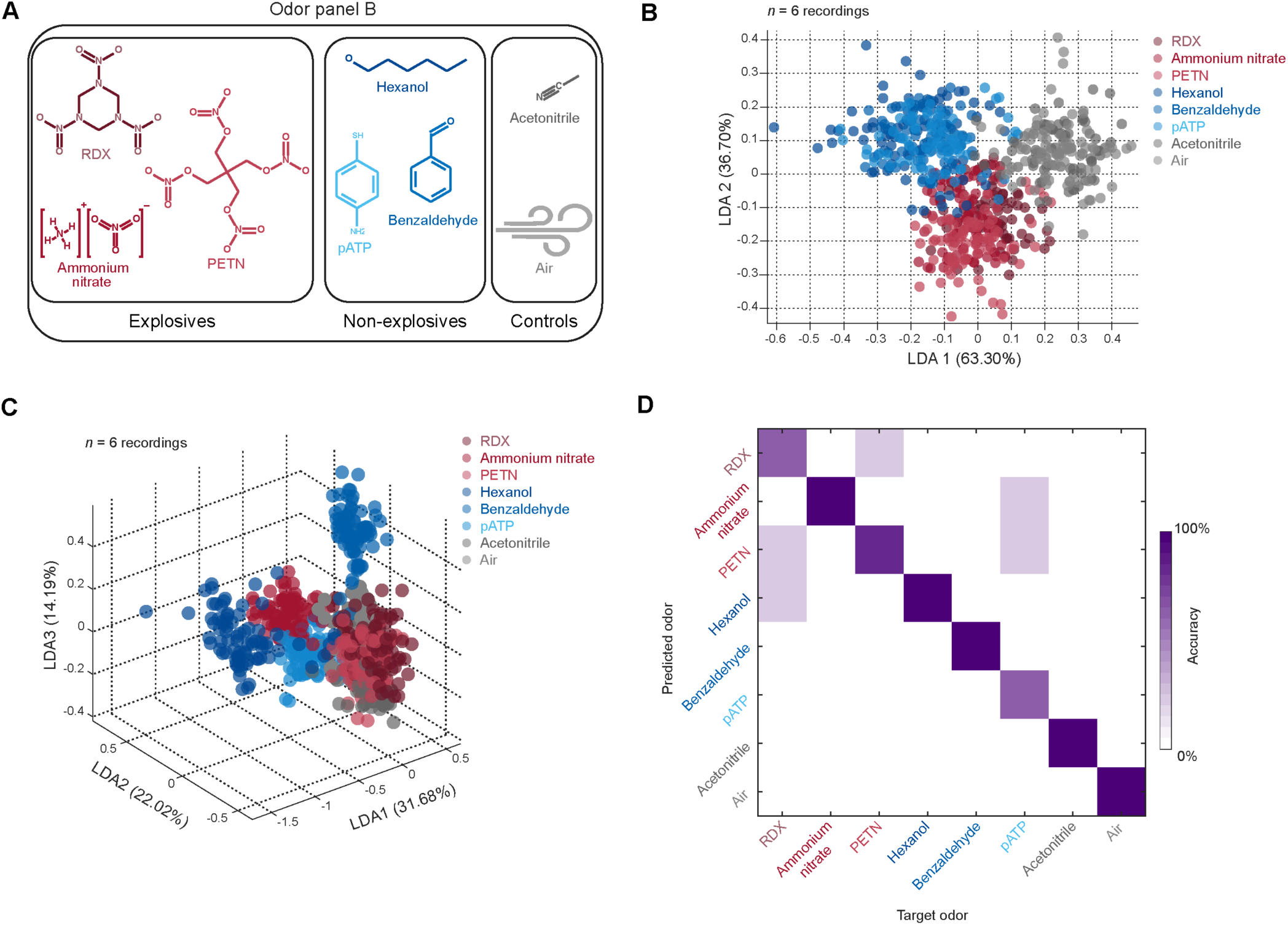
Similar results as in **Figure 4** but for odor panel B. **A)** Odor panel B consisting of RDX (dissolved in acetonitrile), ammonium nitrate (dissolved in mineral oil), PETN (dissolved in acetonitrile), hexanol, benzaldehyde, acetonitrile (control) and air (control). **B)** Visualization of high dimensional data through dimensionality reduction using a 3-class linear discriminant analysis (LDA) is shown. Numbers within parentheses indicate the variance captured along that axis. Chemicals were labelled as explosives (red symbols), non-explosives (blue symbols) or controls (gray symbols). LDA shows clear separation between explosives and non-explosive chemicals for panel B (n = 6 recordings). **C)** Visualization of high dimensional data through a multi class LDA where each chemical was treated as its own class is shown. Numbers within parentheses indicate the variance captured along that axis. These plots show that individual explosives show distinct responses (*n* = 6 recordings for panel B). **D)** Confusion matrices summarizing the results from classification analyses are shown for both odor panels. Note that the matrices are mostly diagonal indicating that both explosive and non-explosive chemicals can be correctly identified based on the neural responses they evoke.

**Supplementary Figure 5:**
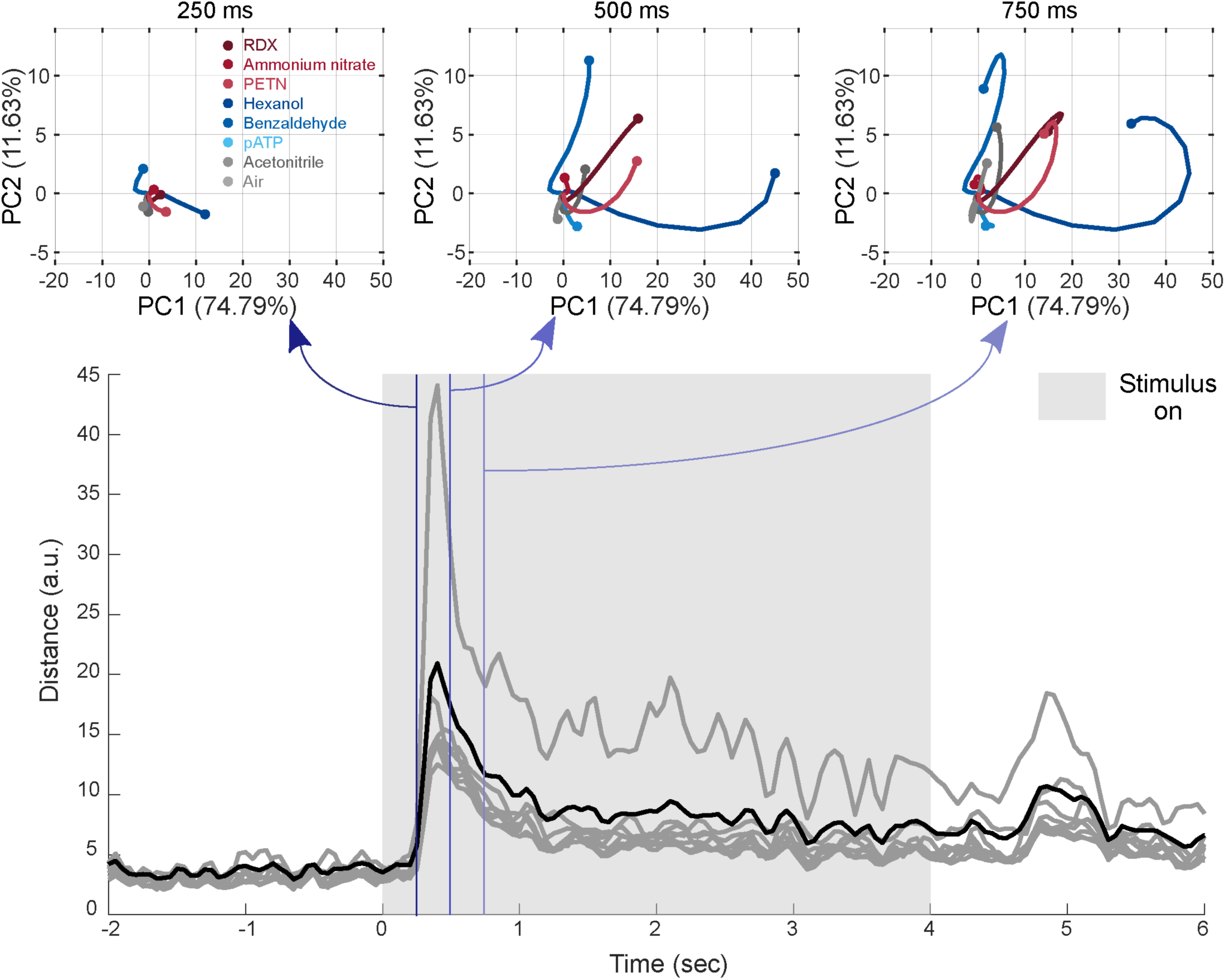
**A)** Rapid identification of chemicals using neural responses: Top panels: high dimensional responses and how they evolve over time are visualized in two dimensions following principal component analysis. Numbers within parentheses indicate the variance captured along that axis. The three panels show evolution of neural responses for the first 250 ms, 500 ms and 750 ms after stimulus presentation. Note that neural responses are similar at 250 ms, but become distinct as they continue to evolve. Maximum separation is reached around 500 ms and the responses start returning towards baseline within 750 ms. Thus, the transient neural responses can be utilized for rapid chemical identification. *Bottom panel*: pairwise distances between neural responses are plotted as a function of time. Black line indicates the mean pairwise distance across all chemicals. Maximum distance indicating maximum separation occurs at ∼350 ms after the onset of chemical stimuli.

**Supplementary Movie 1:** A video showing an actual mobile robotic experiment carried out in the odor localization box. Note that the locust is placed within a custom-made miniature faraday cage and associated amplifiers and wireless data transmitters are placed on the robot platform. Video has been slowed down for illustration purposes.

